# Failing to account for RNA quantity inflates background and leads to the misleading appearance that PRC2 and GFP bind to RNA *in vivo*

**DOI:** 10.1101/2024.11.02.621417

**Authors:** Jimmy K. Guo, Mario R. Blanco, Mitchell Guttman

**Affiliations:** Division of Biology and Biological Engineering, California Institute of Technology, Pasadena, CA 91125, USA; Keck School of Medicine, University of Southern California, Los Angeles, CA 90089, USA

## Abstract

We recently published biochemical and quantitative evidence that challenges the widespread claims that PRC2 binds directly to many RNAs *in vivo*. A recent preprint performs a re-analysis of some of our data using a different metric and claims to identify contradictory results – that PRC2 binds to more RNAs than the well-characterized PTBP1 and SAF-A RNA-binding proteins. The preprint also reports other counterintuitive observations, including that PRC2, and all proteins examined, bind to RNAs that do *not* exist within the same cell and therefore could not interact *in vivo* nor could they be UV-crosslinked. Because these observations are incongruent with orthogonal, non-PCR based, measurements as well as fundamental RNA biology, we explored why their analysis would lead to these unexpected observations. After carefully examining the methods used and regenerating the reported results, we found a very simple explanation for these discrepancies: the authors effectively ignore the vast majority of RNAs in the sample. Here we show that this failure to properly account for the total RNA in each sample leads to inaccurate estimates of the true proportions of each RNA in the sample and therefore the enrichment calculated. We find that the number of RNA reads discarded is dependent on the RNA binding properties of the protein and therefore dramatically inflates background signals (e.g., RNA detected with GFP) while deflating specific signals (e.g., RNA detected with PTBP1). Importantly, we show that this issue alone explains the discrepancy between their observations and our previously reported results. The fact that PRC2 binding can only be observed when analyzed in this way further reinforces the fundamental point of our original paper: the existing evidence in support of direct PRC2-RNA binding requires critical re-evaluation.

## OVERVIEW

We recently published biochemical and quantitative evidence that challenges widespread claims that PRC2 binds directly to many RNAs *in vivo*^1^. Briefly, several observations suggest that many of the previously reported PRC2-RNA interactions may not occur *in vivo*^2–8^. Based on these observations, and because PRC2 has been reported to bind promiscuously to many RNAs^9–11^, we designed an experiment to unambiguously identify non-specific RNA associations that could not have occurred within the cell. Specifically, we mixed UV-crosslinked cells expressing a tagged version of our protein (+tag) with UV-crosslinked cells of a different species that did *<u>not</u>* express the tagged protein (−tag). We purified the tagged-protein and sequenced associated RNAs. In this system, any sequencing reads that align to the species that did not contain the tagged protein (−tag RNA) *<u>must</u>* represent background that:

1. **<u>could not occur *in vivo*</u>** because these RNAs are not in the same cell as the protein, and
2. **<u>could not be UV-crosslinked</u>** because these samples are mixed after crosslinking.

Using denaturing purifications, we observed no detectable RNA enrichment in the −tag samples for any protein^N1^. However, in the case of PRC2, we also observed no detectable RNA enrichment in the +tag samples despite observing robust enrichment for well-characterized RNA binding proteins such as PTBP1 and SAF-A (also known as hnRNPU)^1^. To confirm these observations using direct (non-PCR based) methods, we purified individual PRC2 components from UV-crosslinked cells and visualized the captured RNAs using ^32^P labeling or infrared dyes and observed minimal or undetectable levels of RNA^1^. Indeed, the levels of detected RNA were comparable to those observed upon purification of GFP, a protein that would not be expected to bind to RNA in mammalian cells. In contrast, purification of PTBP1 and SAF-A led to robust levels of RNA detection. Based on these and other results, we “*<u>argue for a critical re-evaluation of the idea that PRC2 and other chromatin regulators bind directly to many RNAs and that such binding is essential for their function”</u>*^1^.

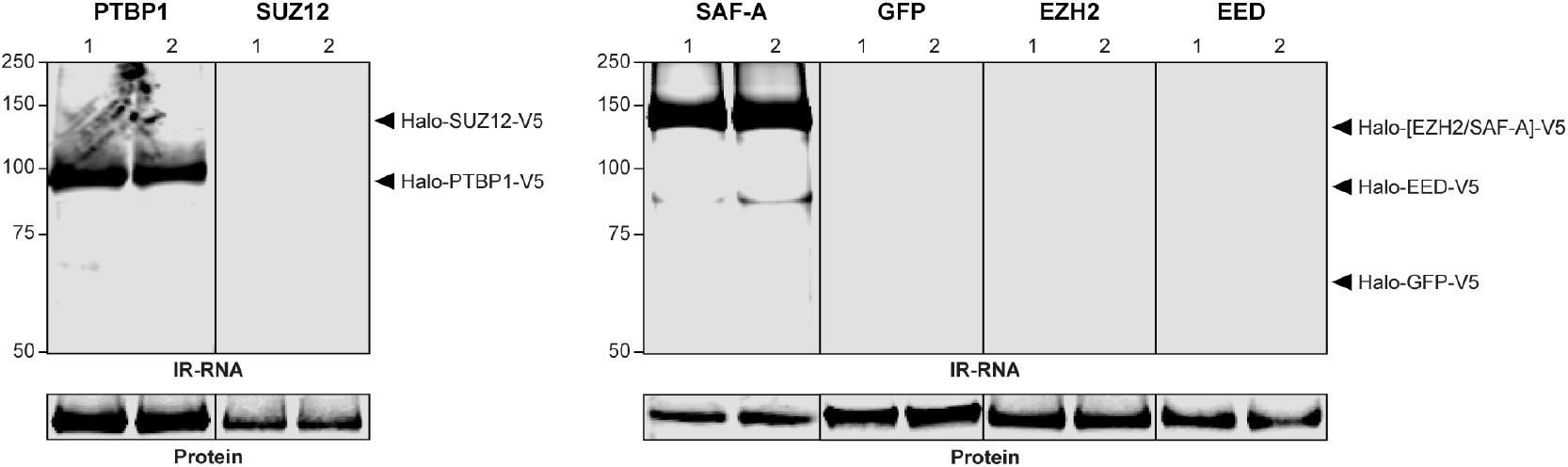

**Figure adapted from Figure S19A in Guo *et al***., **2024**^1^. **See also Figures 4A and S7A**.

Recently, Lee *et al*.^12^ posted a preprint where they re-analyze our sequencing data using a distinct metric and claim to identify a different result. Because the authors were unable to reproduce our results, they claim that this discrepancy must be because we processed the PRC2 samples differently from the other samples. This is simply not the case – all samples were processed identically. Rather, as the authors explicitly state in their preprint, they <u>“</u>*<u>did</u>* ***<u>not</u>*** *<u>attempt to recreate the detailed bioinformatic pipeline</u>*” described in our paper. Instead, they used an entirely distinct metric that does not account for the dramatically reduced RNA amounts observed in these samples.

Numerous observations reported in their preprint highlight that their metric is problematic and leads to results that are incongruent with orthogonal, non-PCR based, measurements as well as fundamental RNA biology. For example, using their metric^12^, they report that:

- <u>**PRC2 binds RNAs that do not exist in the same cell and are not UV-crosslinked**</u> (−tag).
- <u>**PRC2 binds to more RNAs than PTBP1 and SAF-A**</u> (canonical RNA-binding proteins).
- <u>**PTBP1 binds more RNAs not in the same cell** (−tag) **than RNAs that are** (+tag)</u>.

We explored why their analysis would lead to such counterintuitive observations. After careful examination of their methods and regeneration of their reported results, we found a very simple explanation for the differences observed: the authors fail to properly account for the vast majority of RNAs present in the samples. This seemingly simple difference has profound effects that lead to inaccurate results by dramatically inflating the enrichment values *<u>specifically</u>* for background signals (e.g., GFP binding) while deflating the enrichment values of specific signals (e.g., PTBP1 binding). This is precisely why we emphasized the importance of accounting for total RNA amounts when analyzing these data in our paper^1^, which we now explain in more detail below. Rather than contradicting our conclusions, this reanalysis underscores the *<u>exact</u>* point of our paper — there is a need to critically re-evaluate the evidence that PRC2 binds directly to many RNAs *in vivo*.

## RESULTS

### Failure to account for RNA composition leads to inaccurate enrichment calculations that are highly skewed between samples

To explain the issues that lead to this discrepancy, we will briefly describe what we are measuring in an enrichment experiment and what the score represents. Because measurement of RNA binding in a sequencing experiment relies on PCR amplification, which will amplify even small amounts of RNA, the presence of sequencing reads aligning to an RNA alone does not indicate binding^2,13^. Instead, we are asking whether a specific RNA increases in frequency upon capture of a specific protein relative to the frequency of the same RNA in the initial population (“input” sample). To calculate enrichment, we compute the frequency of each RNA in the capture and initial population as the number of reads mapping to the specific RNA divided by the total reads in each sample. The enrichment score for each RNA is the ratio of the frequency observed in the capture divided by the frequency observed in the input. Therefore, an enrichment score >1 means the frequency *<u>increases</u>* in the capture sample relative to the input samples while an enrichment score <1 means the frequency *<u>decreases</u>* in the capture relative to the input.

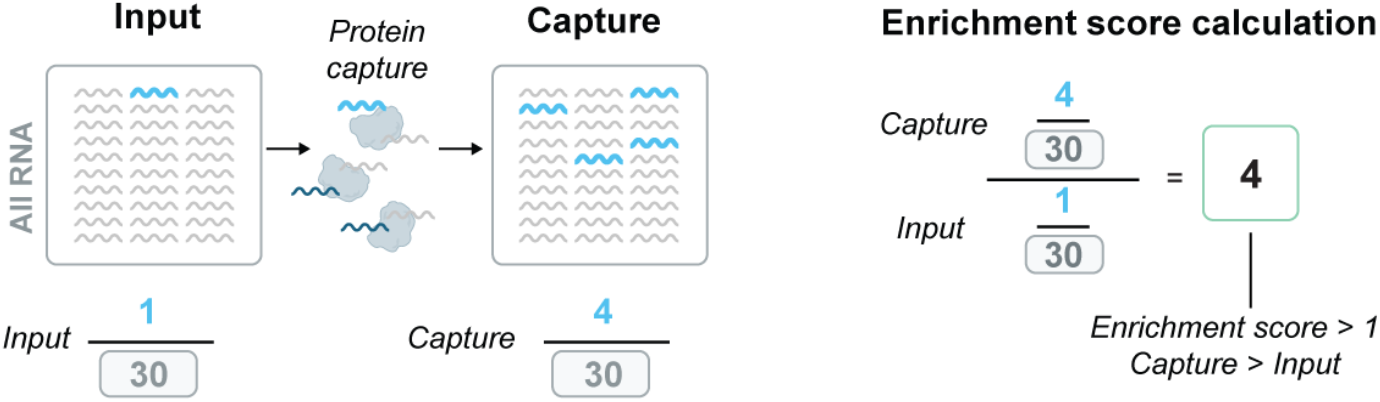

In their preprint, Lee *et al*.^12^ focus only on RNA sequencing reads that they could map uniquely to the genome and exclude the remainder of the reads in the sample from their calculation (the denominator). However, this choice leads to the exclusion of the *<u>vast majority of RNA</u>* present within the sample. To understand why this is, here is a breakdown of the read alignment proportions for one of the input samples.

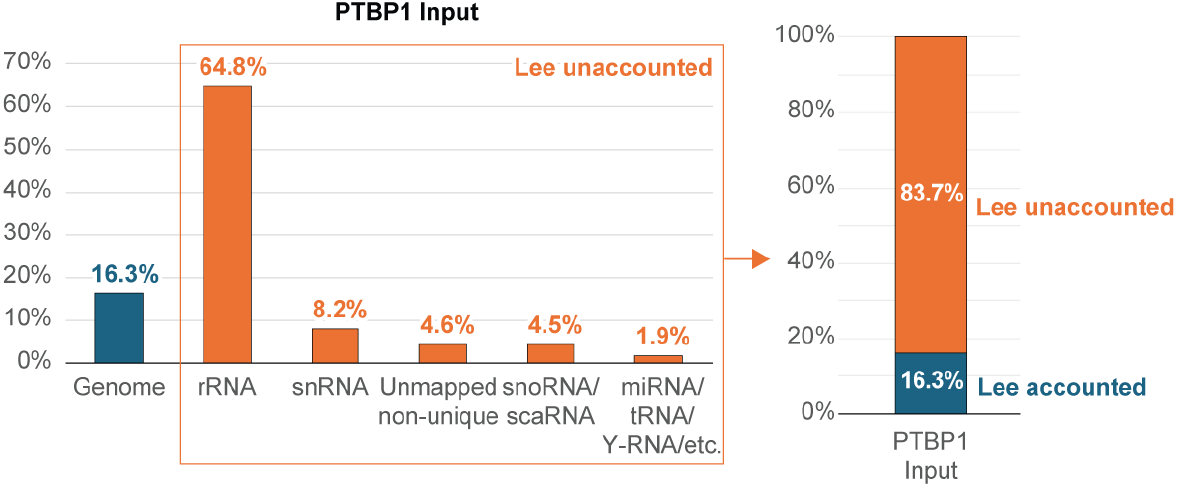

In this case, Lee *et al*.’s analysis^12^ excludes >80% of the RNA reads in the sample. The reads that are excluded correspond predominately to abundant RNA species such as ribosomal RNAs, snRNAs, snoRNAs, tRNAs, etc. The authors rationalize this exclusion due to the reads’ ‘non-unique alignment’^12^. Importantly, the excluded reads are not “non-unique alignments”; they are unique alignments to abundantly expressed RNAs that simply do not align uniquely to the reference genome. While this might represent a bioinformatic distinction, there is *<u>no biologically meaningful reason</u>* that these RNAs should be excluded from measuring the total RNA population present within the experiment.

This distinction is extremely important because enrichment of a given RNA is directly related to the proportion of reads observed relative to the total population of *<u>all</u>* RNAs. Therefore, how one defines this “total” population dramatically impacts the proportion calculated and the enrichment score. Consider an example where the capture and input samples have the same total amount of RNA but a 2.5-fold difference in the number of “genome-mapped” RNAs. If a specific RNA was observed to have an identical number of reads in both capture and input, the RNA is present at the same proportions in both samples (enrichment = 1). However, if we only compute this proportion based on the genome-mapped counts, we would incorrectly conclude that the specific RNA was represented 2.5x more frequently in the capture versus the input sample (enrichment = 2.5).

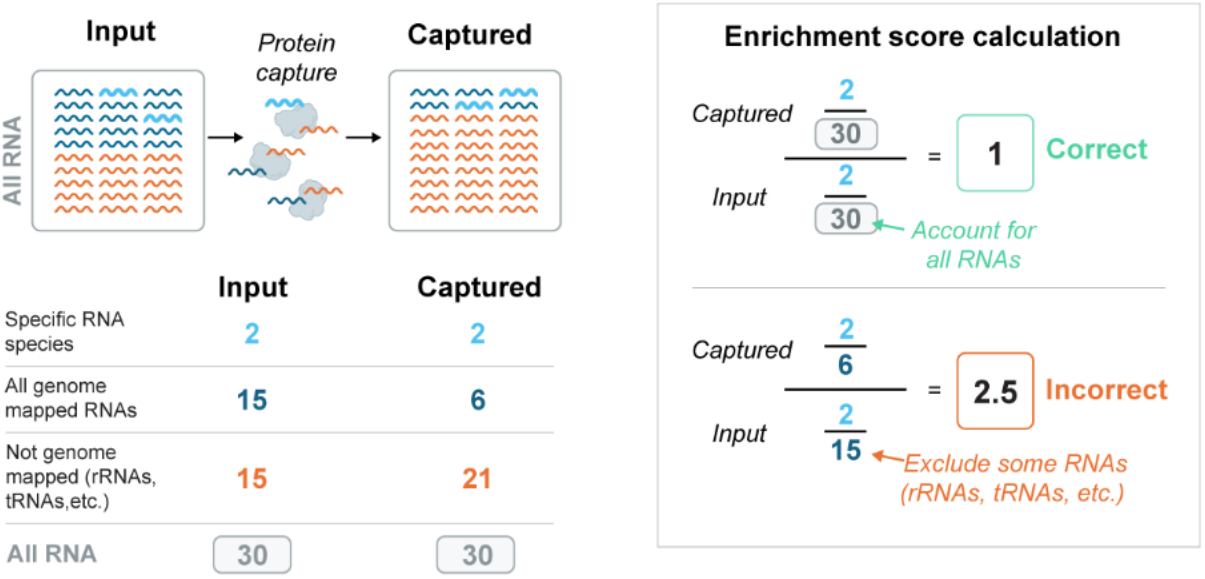

This example is not simply a theoretical one, rather, we see precisely these differences between the compositions of total and genome-mapped reads across the proteins detected. Specifically, the proportion of “genome-mapped” reads relative to the total is not the same between samples. In other words, the number of reads unaccounted for by Lee *et al*.’s analysis^12^ varies *<u>dramatically</u>* between samples corresponding to 2% for EED and 45% for PTBP1, <u>**a >18-fold difference**</u> between these samples. In contrast, we do not observe these fluctuations in the different input samples.

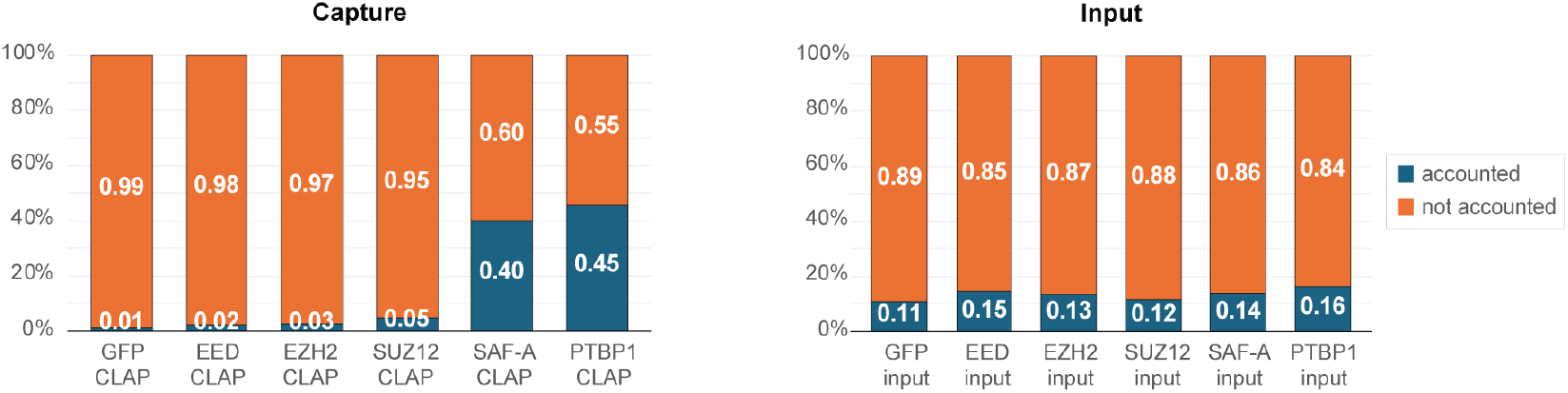

Therefore, focusing only on genome-mapped reads and ignoring the remainder of the RNAs in the sample would lead to skewed estimates of enrichment relative to their actual proportions within the experiment. These “skews” impact different samples differently depending on the proportion of genome-mapped reads within each sample, which varies from protein to protein.

### Exclusion of non-genome-mapped RNAs <u>*preferentially*</u> inflates enrichment values for samples most depleted for RNA binding

Importantly, this variability in the genome-mapped frequency between samples is not random, but rather reflects a *<u>fundamental</u>* property of the RNA binding of the protein measured. Specifically, capture of a protein that binds to RNA regions that can be mapped to the genome, such as intronic regions of pre-mRNAs, might show a higher proportion of genome-mapped reads than input^N2^.

Conversely, a protein that fails to bind to specific RNAs would be expected to show a decrease in the proportion of genome-mapped RNAs and an increase in the frequency of abundant RNAs in the background. (While proteins that bind specifically to abundant RNAs that do not map to the genome, such as ribosomal RNAs, would also lead to a decrease in the proportion of genome-mapped reads, such cases can be reliably detected^14,15^. However, this is not what we observe for PRC2, nor would it make biological sense for this to be the case here.)

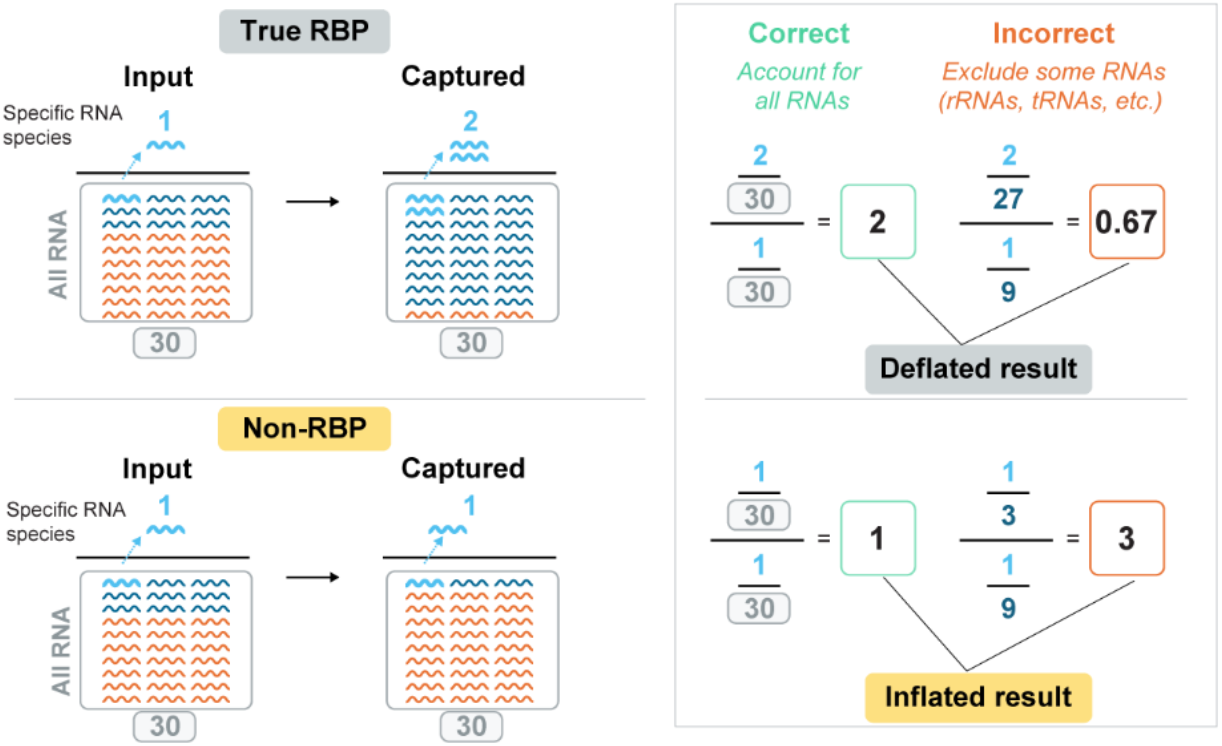

Comparing these proportions between capture and input for different proteins, we observe clear differences in enrichments. For SAF-A and PTBP1, two well-characterized RNA-binding proteins that bind to many specific genome-mapped RNAs, we observe that the proportion of genome-mapped reads increases relative to input; in contrast, for PRC2 and GFP, the proportion decreases relative to input.

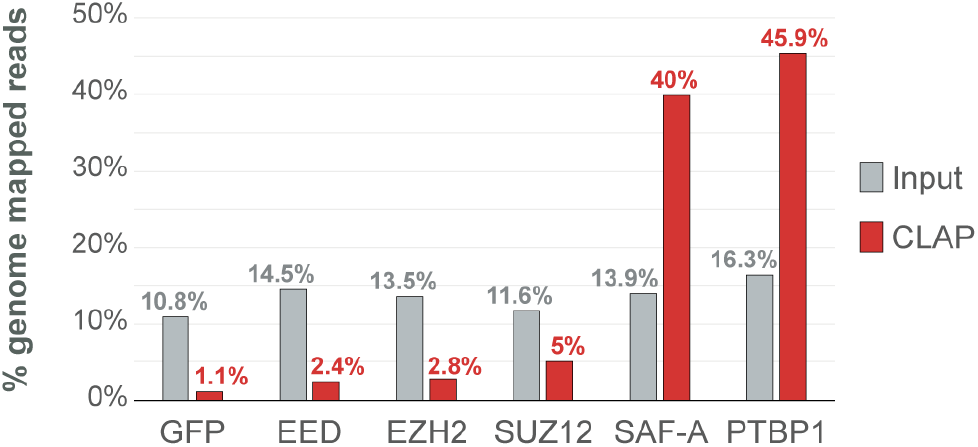

In this way, failing to properly account for all RNAs not only leads to an inaccurate estimate of the true enrichment proportion but *<u>preferentially and dramatically inflates</u>* the enrichment of samples that are most depleted for binding. Notably, in all cases, these dramatically inflated values correspond precisely to the samples that display the lowest amount of detectable RNA when measured by orthogonal non-PCR based measurements.

### The failure to account for RNA composition alone explains the reported discrepancies

This failure to properly account for all RNAs in the sample *<u>alone</u>*^N3^ explains the discrepancy in what Lee *et al*.^12^ report compared to the results in our paper^1^. Because PRC2 and other negative controls like GFP and −tag are the samples that are most depleted for RNA binding and yield the lowest absolute amounts of RNA, these are the samples where the total background population is most likely to represent abundant background RNA sequences. Therefore, it is precisely these samples where artificially “inflating” low RNA yields would be most problematic.

Indeed, computing the enrichment metric using only genome-mapped reads (as Lee *et al*.^12^ does) across all RNAs highlights the problematic nature of this metric. Specifically, one would identify virtually all RNA regions (>80%) as enriched for PRC2 and GFP. It is therefore not surprising that one would also observe the appearance of enrichment over XIST as they report. However, as can be seen, these enrichment values are dramatically inflated across all RNA regions relative to their true proportions within the sample.

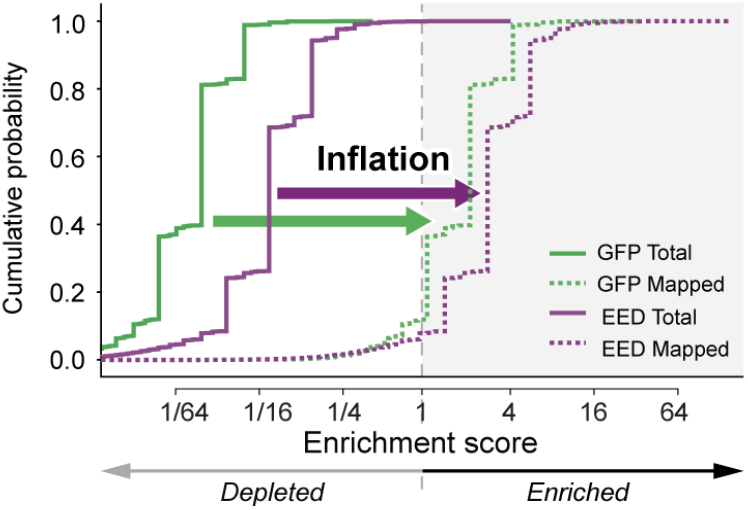

Moreover, we observe the opposite effect for PTBP1: the reported enrichment values are deflated across all RNAs. While some regions are still enriched using this metric, most of the PTBP1 binding sites are not detected. Accordingly, this deflation of the positive control PTBP1 and inflation of the negative control GFP leads to the erroneous appearance that GFP binds to more RNAs than PTBP1.

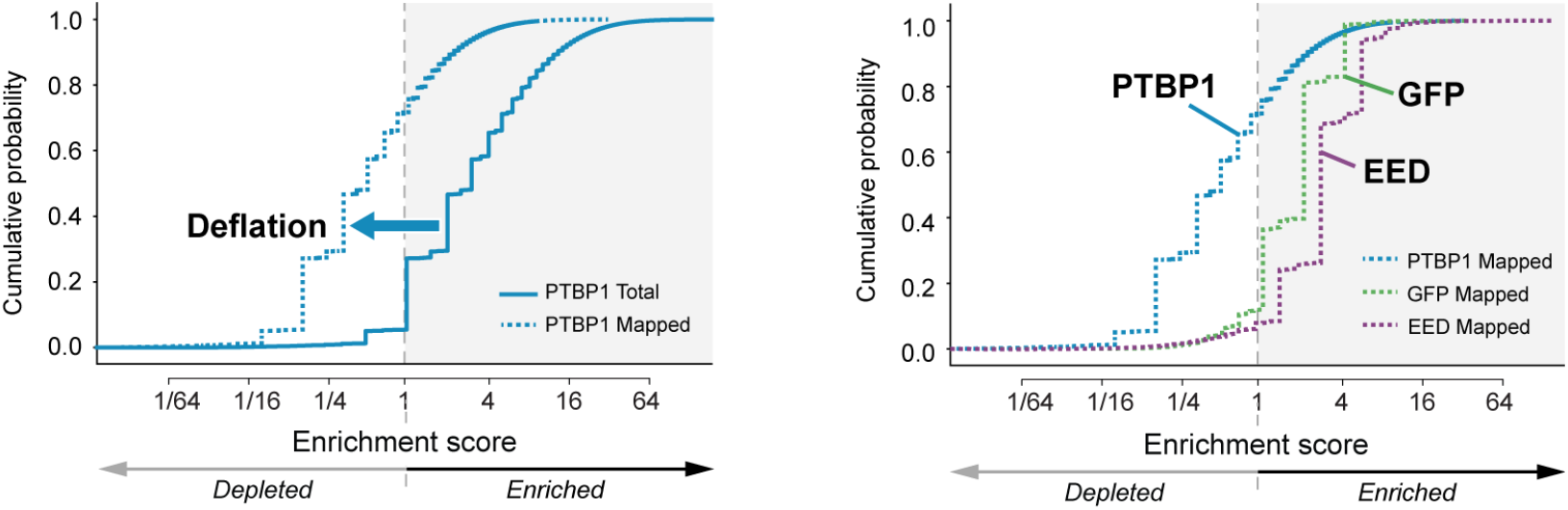

It is precisely this failure to account for the vast majority of RNAs in the sample and the resulting background inflation that leads to the other counterintuitive observations reported^12^.

For example:

- They report that PRC2 binds to *<u>more</u>* RNAs than PTBP1. This observation is incompatible with the massive difference in RNA amounts observed in both ^32^P and IR-gels as well as the higher crosslinking efficiency^16^ and *in vitro* binding affinity for PTBP1^1^. This observation occurs not because PRC2 binds to more RNAs but rather because their metric preferentially inflates PRC2 values *<u>and</u>* deflates PTBP1 values precisely because of their striking differences in binding.
- They report *<u>stronger</u>* enrichment on more RNAs for PTBP1 on RNAs that are *<u>not</u>* expressed within the same cell (−tag) than on RNAs that are expressed in the same cell (+tag). By its very definition, this cannot be true. Rather, this observation occurs because they are specifically inflating the enrichment of the −tag sample precisely because the protein does not bind to RNA in these samples while deflating the enrichment of the +tag sample precisely because it does bind.
- They report that *<u>all proteins</u>* are enriched for binding in the −tag samples. This reflects the fact that their metric specifically inflates these enrichments precisely because the protein does not bind to RNA in these samples.
- They report that they see *<u>no difference</u>* in enrichment between CLIP and CLAP samples, an observation that is incompatible with orthogonal PCR-independent measurements presented in our paper^1^. Their observation occurs because the proportion of genome-mapped reads decrease in CLAP relative to CLIP for PRC2 and GFP, leading to inflation of their enrichments. By contrast, these proportions do not meaningfully change for PTBP1 or SAF-A and therefore do not show a difference (as we reported) between CLIP and CLAP.
- They report that SAF-A is *<u>not</u>* enriched on Xist^12^ even though it is a well-established Xist binder with clear functional roles in Xist-mediated silencing^4,17–19^. Conversely, they report that all PRC2 components are enriched on Xist, even though numerous studies that purified Xist and characterized its binding proteins (including from J.T. Lee’s own lab^5^) have consistently failed to detect such an interaction with *<u>any</u>* of these proteins. This observation occurs because their metric preferentially deflates values for SAF-A and inflates values for PRC2 precisely because of their clear differences in binding.

## CONCLUSION

Given the issues discussed above, it is important to note that the authors do not provide support for the validity of their metric. In fact, the only “validation” presented is their claim that they have replicated our results for everything except PRC2^12^. However, this is simply not true. Rather, what they meant was that they *<u>also</u>* observe enrichment for PTBP1 and SAF-A. However, this observation does not indicate that their metric is meaningful, but simply highlights that these proteins are strong enough binders that they can be observed even when their values are deflated. In contrast, PRC2 enrichment can *<u>only</u>* be observed when analyzing the data using a problematic and unvalidated metric – a metric on which previous claims of PRC2 binding are similarly based^20^.

Although Lee *et al*. claims that our analysis is ‘unconventional’^12^, there is nothing unconventional about accounting for sample abundance in sequencing data. To the contrary, numerous studies have highlighted how failing to account for sample abundance can lead to inaccurate and misleading conclusions for many different applications, from measuring gene expression by RNA-Seq^21^ to mapping protein binding to genomic DNA using ChIP-Seq^22,23^. The issues with previous (‘conventional’) approaches and why they can lead to incorrect interpretations *<u>specifically</u>* when studying proteins that do *<u>not</u>* bind to RNA (e.g., GFP, −tag) is precisely the point made in our paper^N4^.

These technical issues aside, the claim that PRC2 binds to −tag RNAs, which by its very experimental design could *<u>not</u>* occur, clearly highlights the <u>***exact***</u> point of our paper. While one could argue about what metric is most appropriate, this is a distraction from the fundamental scientific issue at the heart of this debate. There are numerous functional and biochemical reasons to question whether PRC2 binds to RNA *in vivo* (discussed in Guo *et al*.^24^). Whether one analyzes the data as we believe is accurate and observe global depletion of PRC2 binding to RNA, or whether one analyzes it in a different way and observe global enrichment of PRC2 on RNAs that could <u>***not***</u> bind *in vivo*, both lead to the same fundamental conclusion stated in our paper: there is clearly a need to rigorously and critically re-evaluate the existing evidence supporting the claim that PRC2 binds directly to RNA *in vivo*^N5^.

## METHODS

### Datasets

All CLAP and input datasets used are available in the Gene Expression Omnibus (https://www.ncbi.nlm.nih.gov/geo/) under accession number: GSE253477.

### Data analysis

The complete analysis pipeline and all associated files required to perform the analyses described are available at https://github.com/GuttmanLab/CLAPAnalysis.

### Read processing and alignment

Paired-end RNA sequencing reads were trimmed to remove adaptor sequences using Trim Galore! v0.6.2 and assessed with FastQC v0.11.8. Read pairs were then aligned to a combined genome reference containing the sequences of repetitive and structural RNAs (ribosomal RNAs, snRNAs, snoRNAs, 45S pre-rRNAs, tRNAs, and others) using Bowtie2. The genome reference file containing these repetitive and structural RNA sequences are available at https://github.com/GuttmanLab/CLAPAnalysis. The remaining reads were then aligned to a combined genome reference containing the mouse (mm10) and human (hg38) genomes using STAR aligner^25^. PCR duplicates were removed using the Picard MarkDuplicates function. For human and mouse mixing experiments (e.g., +tag/-tag experiments), only reads that mapped uniquely in the genome and unambiguously to the human or mouse genomes were kept for further analysis. For experiments done in a single species, the appropriate reference genome and alignments were used (mm10 for mouse and hg38 for human).

### Gene window enrichment calculations

All human (hg38) and mouse (mm10) annotated genes (RefSeq, downloaded from UCSC GRCh38 and GRCm38, respectively) were used as a reference set except for the genes encoding the transfected proteins. We treated exonic regions and intronic regions of each annotated gene as separate reference genes for computing enrichment. For each reference gene, we enumerated 100 nucleotide windows that span across the gene; for each window, we calculated: (i) the number of reads overlapping the window in the protein elution sample (e.g., CLAP) and (ii) the maximum of either the number of observed reads over the window or the median read count over all windows within the gene in the input sample. Because all windows overlapping a gene should have the same expression level in the input sample, this approach provides a conservative estimation of the input coverage because it prevents windows from being scored as enriched if the input values over a given window are artificially low due to stochastic fluctuation, while at the same time accounting for any non-random issues that lead to increases in read counts over a given window (i.e., alignment artifacts leading to non-random assignment or pileups).

To directly compare the number of reads within each window between sample and input, we normalized each window count by the total number of reads sequenced and the overall complexity within each sample. For example, if one sample was sequenced twice as deeply as another, then we would expect to observe – on average – twice as many reads over a given window for that sample. To account for the differences in overall complexity between samples, we scaled the total number of sequenced reads by the proportion of genome-mapped reads within each sample.

For each window, we computed enrichment by dividing the normalized sample counts by the normalized input counts. Nominal *p*-values were calculated for each window using a binomial test where *k* (number of successes) is defined as the number of reads in the protein elution samples within the window, N (number of trials) is the sum of the number of reads in the protein elution and input samples, and *p* (probability of success) is the expected number of reads in the elution sample divided by the sum of the expected number of reads per window in elution and input samples. (The expected number of reads is defined as the total number of reads scaled by the proportion of aligned reads within each sample).

### Plotting and visualization

Bar plots depicting distribution of aligned RNAs were plotted using the alignments generated as described above. Specifically, “genome-mapped” RNAs refer to reads aligning uniquely to mouse (mm10) or human (hg38) genomes, whereas “non-genome-mapped” RNAs refer to either reads aligning to repetitive and structural RNAs (ribosomal RNAs, snRNAs, snoRNAs, 45S pre-rRNAs, tRNAs) or unaligned reads. Cumulative distribution plots were generated using 100 nucleotide windows as computed above, using either “genome-mapped” reads only or all reads.

## Footnotes

N1 Lee *et al*.^12^ claims that our paper lacked certain “controls”, such as −tag samples for PTBP1 and SAF-A CLAP. In fact, these were generated and show >99% depletion of −tag RNAs for all proteins studied. We simply chose not to highlight these samples because they had no direct relevance to the results presented. Specifically, the −tag experiment was used to determine whether CLIP of PRC2 detects binding when it could not occur and whether CLAP could eliminate this. While the same is true for PTBP1 and SAF-A, these experiments were performed for a distinct purpose to confirm that CLAP does not impact the detection of known interactions. Quantifying binding in the +tag samples does not rely on the −tag samples because samples displaying low +tag *and* low −tag signals are *<u>not equivalent</u>* to samples displaying high +tag *and* high −tag signals (computing the +tag:−tag ratio would distort this distinction).

N2 While the relative proportion of genome-mapped to total reads *<u>alone</u>* does not indicate whether a protein binds to a specific RNA, we use this example to highlight why it is important to account for global shifts to accurately measure enrichment on specific RNAs. We discuss other important considerations for defining specific binding sites in our paper^1^ and in other published work^14,15^.

N3 While Lee *et al*.^12^ enumerates a list of other potential differences, including the window sizes used for the analysis, *p*-value cutoffs chosen, input samples used for normalization, or the statistical distribution used, none of these can explain the discrepancies they described. Briefly, while there are certainly some arbitrary choices that are required for all analyses, our results are robust to these precise parameters. For example, our results are not dependent on 100-nt windows. We see similar effects for other window sizes and the specific IGV plots shown throughout the paper were generated using single nucleotide windows (see STAR Methods^1^, “Plotting and visualization”). Furthermore, our results are robust to the precise *p*-value cutoff chosen; we show the results of different cutoffs in our paper^1^ (see Figure S7C). While we chose to merge the input samples from experiments performed in the same cell-types (which have the same expression) to reduce stochastic variation, we observe the *<u>same</u>* results when normalizing to their *<u>paired</u>* input samples. Finally, the effects are certainly not due to the choice of a Binomial distribution instead of a Poisson distribution since the Poisson distribution is simply an estimator of the Binomial distribution when *n* (number of reads over a window) is large, an assumption that is not generally true in these data (a brief explanation can be found <u>here</u>). While there are other important differences in how we compute enrichment compared to Lee *et al*., they are not the reason for the large discrepancy reported by Lee *et al*.^12^.

N4 Specifically, we state in the main text of our paper^1^: *“Despite these issues, the CLIP procedure itself is not the problem; rather, complications can arise from how the data are interpreted.”* “These quantitative differences may explain some of the apparent discrepancies between conclusions from previous CLIP studies, including more stringent variants such as denaturing CLIP (dCLIP), and those reported here.” These *“non-random associations that often show discrete UV-dependent and protein-specific “peaks” that could be mistaken for legitimate binding sites using standard analytical methods.”*

N5 We encourage those interested in exploring further to read our paper, explore our data (as well as other previously published datasets), and draw their own conclusions. All data and analysis pipelines are available at: https://www.ncbi.nlm.nih.gov/geo/query/acc.cgi?acc=GSE253477 (GEO: GSE253477) and https://github.com/GuttmanLab/CLAPAnalysis.

